# Getting the gist faster: Blurry images enhance the early temporal similarity between neural signals and convolutional neural networks

**DOI:** 10.1101/2021.08.22.451834

**Authors:** David A. Tovar, Tijl Grootswagers, James Jun, Oakyoon Cha, Randolph Blake, Mark T. Wallace

## Abstract

Humans are able to recognize objects under a variety of noisy conditions, so models of the human visual system must account for how this feat is accomplished. In this study, we investigated how image perturbations, specifically reducing images to their low spatial frequency (LSF) components, affected correspondence between convolutional neural networks (CNNs) and brain signals recorded using magnetoencephalography (MEG). Using the high temporal resolution of MEG, we found that CNN-Brain correspondence for deeper and more complex layers across CNN architectures emerged earlier for LSF images than for their unfiltered broadband counterparts. The early emergence of LSF components is consistent with the coarse-to-fine theoretical framework for visual image processing, but surprisingly shows that LSF signals from images are more prominent when high spatial frequencies are removed. In addition, we decomposed MEG signals into oscillatory components and found correspondence varied based on frequency bands, painting a full picture of how CNN-Brain correspondence varies with time, frequency, and MEG sensor locations. Finally, we varied image properties of CNN training sets, and found marked changes in CNN processing dynamics and correspondence to brain activity. In sum, we show that image perturbations affect CNN-Brain correspondence in unexpected ways, as well as provide a rich methodological framework for assessing CNN-Brain correspondence across space, time, and frequency.

## Introduction

The human visual system has been characterized as a hierarchical system that begins with extraction of information about simple features (e.g., oriented contours) registered by neurons whose receptive fields are retinotopically organized, followed by increasingly refined analysis of more complex aspects of the visual scene via neurons with increasingly large receptive fields (Hubel & Wiesel, 1977; Maximilian Riesenhuber, 1999; Serre, Oliva, & Poggio, 2007; Vinckier et al., 2007). Generally, inspired by this biological organization, convolutional neural networks (CNNs) built for image classification have been similarly constructed such that early convolutional layers register simple features in small receptive fields, followed by pooling layers that progressively increase receptive field size, allowing subsequent convolutions to extract complex features that are then passed to fully connected layers for classification (Kietzmann, Mcclure, & Kriegeskorte, 2019; Lecun, Bengio, & Hinton, 2015; Richards et al., 2019). Although neural networks are biologically implausible in some ways, such as weight sharing and backpropagation, they are nevertheless increasingly recognized as useful models of neural processing (Cadieu et al., 2014; Khaligh-Razavi & Kriegeskorte, 2014; Yamins & DiCarlo, 2016). Still, recent studies question the generality of the correspondence between neural networks and neural activity, noting that the relationship between fMRI activation patterns and CNNs is considerably weakened when visual images are degraded or comprised of artificial objects (Xu & Vaziri-Pashkam, 2021). However, it remains possible that the poor temporal resolution of fMRI obscures the category structure that emerges as a function of the temporal dynamics of processing within the visual stream (Carlson, Tovar, Alink, & Kriegeskorte, 2013; Cichy, Pantazis, & Oliva, 2014; Wardle & Baker, 2020). Thus, the degree of correspondence between CNN models and dynamic brain signals associated with degraded visual images remains an open question. In the current work, we address this question by measuring brain responses to degraded images using high temporal resolution magnetoencephalography (MEG) and comparing these to performance in a number of CNNs.

The form of visual image degradation we have focused on is motivated by the coarse-to-fine manner by which the brain is thought to optimize object recognition (Bar, 2003a; Bar, Kassam, Ghuman, Boshuan, et al., 2006; Petras, ten Oever, Jacobs, & Goffaux, 2019). This view posits that low spatial frequency information is processed by the faster magnocellular pathway (Kauffmann, Ramanoël, Guyader, Chauvin, & Peyrin, 2015; Tootell, Silverman, Hamilton, Switkes, & De Valois, 1988), which creates an initial coarse representation of the image/object. Called “scene gist”, those initial representations or “hunches” are then refined as more detailed information emerges in the form of high spatial frequencies processed by the slower parvocellular pathway traveling through the ventral visual stream (Bar, 2003a; Bar, Kassam, Ghuman, Boshuan, et al., 2006; Bruner & Potter, 1964; Snodgrass & Hirshman, 1991; Tootell et al., 1988). Low frequency information was initially thought to enhance processing within the ventral visual stream through feedback signals originating in the orbitofrontal cortex (OFC) to category selective areas in inferotemporal (IT) cortex (Bar, 2003; Bar, Kassam, Ghuman, Boshuan, et al., 2006). However, recent evidence suggests that feedback processes are more diffuse along the ventral visual stream. For example, an fMRI occlusion paradigm that selectively manipulated the spatial frequency along different receptive fields found that low frequency information is conveyed through feedback signals throughout the ventral visual stream, including in primary visual cortex (Revina, Petro, & Muckli, 2018). Additionally, high spatial frequency processing domains are segregated from low spatial frequency processing domains as far upstream as V4, indicating that unique spectral information is preserved within feedforward processing (Lu et al., 2018). Collectively, it thus appears that coarse-to-fine processing comprises a combination of dynamic feedforward and feedback interactions. This implies that the extent to which the brain relies on low spatial frequencies to initiate top-down processes depends on the available spectral and contextual information present in an image.

Modeling the visual system requires capturing the dynamics of object recognition under a variety of task constraints, including degraded images that necessitates varying degrees of feedback/top-down processing. The sluggish fMRI signal makes it difficult to differentiate between the dynamics of early feedforward and later feedback processes; these dynamics take on particular importance as we go beyond assessing CNN-Brain correspondence with natural images (Cichy, Khosla, Pantazis, Torralba, & Oliva, 2016; Güçlü & van Gerven, 2015, 2017; Khaligh-Razavi & Kriegeskorte, 2014; Kietzmann, Spoerer, et al., 2019; Kong, Kaneshiro, Yamins, & Norcia, 2020; Mehrer, Spoerer, Jones, Kriegeskorte, & Kietzmann, 2021; Schrimpf, Kubilius, Hong, Majaj, Rajalingham, Issa, Kar, Bashivan, Prescott-Roy, Schmidt, et al., 2018). Behavioral studies have shown that CNNs differ from human vision in terms of susceptibility to the impact of image distortion on object recognition. For example, distortions such as color remapping, low pass filtering and high pass filtering reduce CNN performance in object classification but have considerably less effects on human performance (Geirhos et al., 2018). Thus, the effect of image perturbations on CNN-Brain correspondence is best suited using brain signals measured with high temporal resolution techniques such as M/EEG.

Consequently, in the current work, we have studied the temporal correspondence between neural activity collected using MEG and a diverse set of CNN architectures for clear images as well as for degraded images containing only low spatial frequency components. The added temporal resolution in the MEG allows us to make inferences regarding how the correspondence between MEG signals and the CNN activations evolves throughout the stimulus presentation, and whether image perturbations change the timing of when the correspondence emerges. We predicted that for all images (clear and degraded) there would be a general temporal relationship between CNN layer depth and the time course of the MEG signal following stimulus presentation. In such a framework, shallow CNN layers (those close to the input layer) will correspond to earlier times in the MEG signal and deep CNN layers will correspond to later times in the MEG signal when participants have been allowed more time to fully process an object. However, for the low spatial frequency images, we hypothesized that an enhanced contribution of top-down feedback would result in the more rapid emergence of correspondence between deep CNN layers and MEG signals.

## Methods

### Data Set

We used a data set originally published in Grootswagers et al., 2017. The data comprised results from 20 participants (four men; mean age = 29.3 years) with normal or corrected-to-normal vision participating in an MEG experiment. Stimuli consisted of 48 grayscale images comprised of an even split of animate and inanimate objects on a phase-scrambled natural image background (Figure 1A). Importantly, the stimuli did not include humans and better accounted for shape and other confounds present in the stimuli in other datasets (Grootswagers & Robinson, 2021). The objects were presented in a clear condition and a degraded condition intermixed within eight blocks, resulting in 32 trials for each respective clear and degraded object. Degraded images were constructed by convolving a sombrero function over a Fourier transformed image and selecting varying radii of pixels from the image, resulting in different degrees of low spatial frequency blur (Figure 1A and Supplemental Figure 1). Given that different types of blurring can affect object recognition to different degrees (Kadar & Benshahar, 2012), each image was blurred based on the results from a separate online MTurk experiment with blur being set as the radii by which at least 25% of participants could name the object in a naming task. Stimuli were projected (at 9° × 9° visual angle) on a black background for 500ms with a random intertrial interval between 1000 and 1200 milliseconds. Participants categorized the stimulus as animate or inanimate as fast and accurately as possible. Motor responses were remapped between alternating blocks to avoid potential motor confounds. Prior to the MEG experiment, a familiarization task was used to make sure that all participants could categorize all clear and degraded stimuli as animate or inanimate with accuracy scores of at least 80%. Each MEG recording was done with a whole-head MEG system (model PQ1160R-N2; KIT, Kanazawa, Japan) while participants lay in a supine position inside a magnetically shielded room. Trials were sliced into 700ms epochs spanning from 100ms prior to stimulus onset to 600ms post stimulus onset.

**Figure 1.**
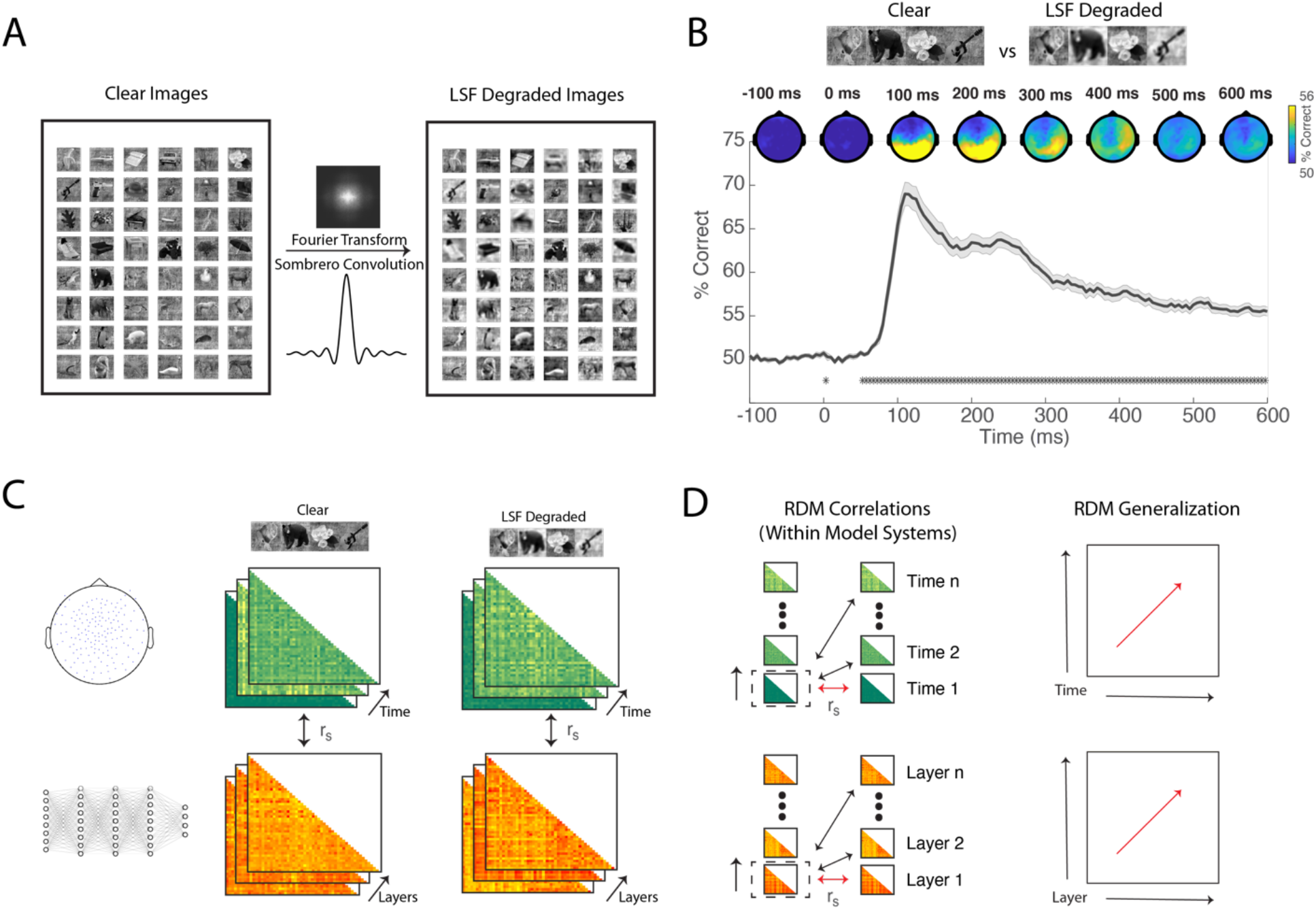
Study Design and Analysis Overview. (A) Stimuli consisted of 48 achromatic visual objects that included 24 animate and 24 inanimate objects shown in prototypical viewpoints. No human faces were included in the data set. Images were placed on a phase scrambled background. Images were degraded using a Fourier transform and sombrero function to preserve the low spatial frequencies individually calibrated for each image to preserve recognition. (B) Time-resolved decoding plot between clear and degraded images for MEG signals. On the x-axis time in milliseconds; on the y-axis decoding performance. Significance is indicated with asterisks above the abscissa using Wilcoxon signed-rank test against chance decoding (50%), FDR corrected, q < 0.025. On the top of the plot, exploratory searchlight analysis shows the topographic distribution of the decoding performance in time. (C) Representational Dissimilarity matrices were calculated using LDA 4-fold cross validation MEG signals and across layers using squared Euclidean distance for each layer activation. RDMs in time and across layers were correlated between MEG and neural networks (D) The evolution of the signal was assessed by correlating RDMs iteratively across all timepoints for MEG signals and layers for neural network activation, creating a RDM generalization matrix.

### Decoding between clear and degraded images

To determine when differences emerged in time between clear and low spatial frequency degraded images (Figure 1B), we trained and tested a classifier using linear discriminant analysis (LDA) (Duda & Hart, 2001). In this procedure, we used a four-fold, leave one-fold out train to test split, iteratively changing which folds were trained and tested. We performed this analysis using all of the MEG sensors to compute an overall decoding classification performance for clear unfiltered images and for low spatial frequency degraded images. Statistical significance was computed by comparing the decoding performance to chance level decoding (50%), correcting for multiple comparisons using FDR correction. In addition, to obtain a topographic estimate of how clear and degraded images are distinguished in the brain, we performed a moving searchlight analysis (Etzel, Zacks, & Braver, 2013), iteratively decoding clear from degraded images at each sensor and its immediate surrounding neighboring sensors. This procedure produced a topographic heat map of decoding performance along 100ms intervals, spanning from 100ms prior to stimulus presentation to 600ms post stimulus presentation.

### Neural RDMs

To capture the time resolved neural relationship between objects, we used representational similarity analysis (RSA) (Kriegeskorte, Mur, & Bandettini, 2008). For each exemplar, we performed pairwise decoding using LDA with four-fold leave one-fold out cross validation for all stimulus comparisons within the clear and low spatial frequency degraded images until we had decoding scores across all possible exemplar comparisons across all time points. Together, these formed time-resolved representational dissimilarity matrices (RDMs) for clear and degraded images respectively (Figure 1C).

### CNN RDMs

Network RDMs were similarly constructed using RSA (Figure 1C). We chose a diverse set of six CNNs of varying depth as well as different types of connections, including skip connections (He, Zhang, Ren, & Sun, 2015), inception layers (Szegedy et al., 2014), and recurrence (Kubilius et al., 2018). Instead of using cross validation, we used the square Euclidian distance between layer activations for each exemplar comparison to build the RDMs. We chose this distance measurement to make the fewest necessary assumptions regarding the relationship between layer activations for each object. Note that in this process, each n × n layer activation is converted to 1 × n vectors preserving the relative relationship of activation within each layer. To measure network dynamics and correspondence with brain activity, we selected all of the convolutional and fully connected layers within each network. However, we also performed the analysis using all possible computations within each network, including pooling layers where convolutional features are pooled, ReLU activation functions that convert all negative values to zero, and normalization layers that scale and center the activations, finding qualitatively similar results.

### Probing Neural and Network Dynamics Separately

To probe whether CNNs and brain activity exhibit similar dynamics when processing clear images and degraded images, we correlated RDM averaged across all participants for each time across all other RDMs in our stimulus window (−100ms to 600ms). This analysis was performed using participant averaged brain RDMs instead of individual RDMs in order to have more stable neural representations. The RDMs are consistently changing in time, so by doing a cross correlation across timepoints we are capturing the dynamics of how each participant processed the clear and degraded objects. We performed a similar procedure separately for CNNs, using layers instead of time (Figure 1D). Given that the neural time window included time before stimulus presentation and that additional time elapses for neural signals to travel from the retina to visual cortex, we chose to begin the cross correlations 50 ms after stimulus. Additionally, since each of the different CNN architectures contains different depths and layer, we interpolated each of the network activations to fit the same dimensions as the brain RDMs (Figure 2A) using a nearest-neighbor interpolation. The nearest-neighbor interpolation duplicates individual pixel values to fit the brain RDM values. We performed this analysis for clear images and for degraded images, and then correlated the relative representational geometry between the clear and degraded images for the brain RDMs and the various CNN architectures separately.

**Figure 2.**
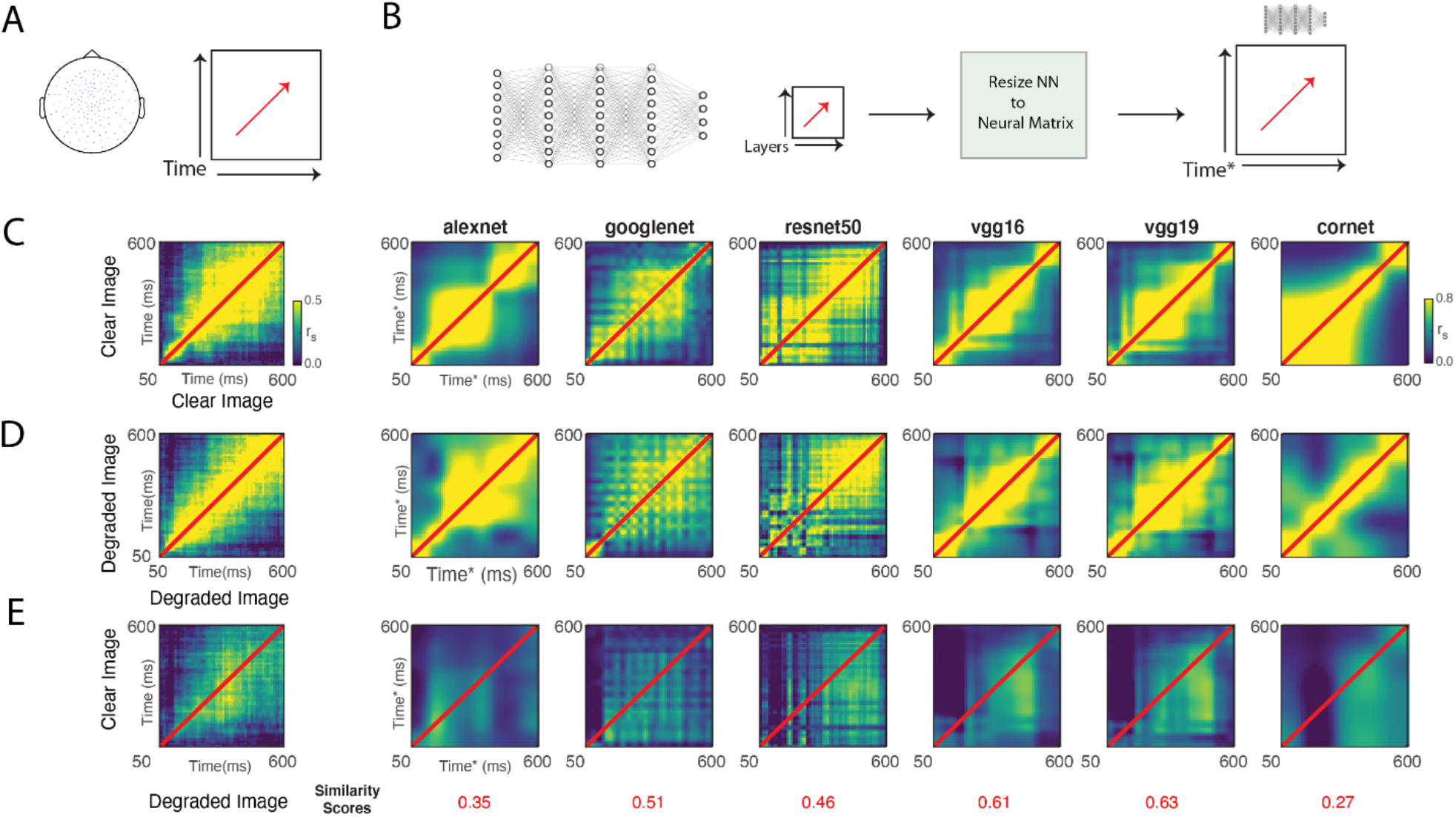
Image processing dynamics to clear and degraded images in brains and CNNs. Correlations between neural RDMs (using all channels) for clear, degraded, and between conditions were calculated using a Spearman correlation. To compare the temporal evolution of the signals in time between neural RDMs and network RDMs, the neural RDM time interval was restricted to approximately when the signals first appear in V1. (B) Neural network RDMs across a wide assortment of networks that include shallow CNNs and deep CNNs, skip connections, as well as recurrence. To compare to the neural RDMs, the RDMs of the various networks were scaled to match the dimensions of the neural RDM using a nearest neighbor interpolation. (C-E) Resulting RDM correlation matrices with neural RDMs on the leftmost column and CNN RDMS to the right for clear images (C) degraded images (D), and the cross correspondence between clear and degraded images (E). For panel E, the similarity score was calculated as (1-squared Euclidean distance) between neural RDMs and CNN RDMs shown on the abscissa.

### Correspondence between brain and CNN RDMs

To relate brain RDMs to the CNN layer specific RDMs, we used a non-parametric Spearman correlation between the brain and CNN matrices across each time point and network layer to avoid making any assumptions of linearity for the Brain-CNN correspondence. We then measured the time in which each CNN layer was maximally correlated to brain data. In addition, we calculated the lower bound of the brain noise ceiling for clear image presentations and degraded image presentations separately. The lower bound of the noise ceiling was approximated by iteratively calculating across all participants the mean correlation between each individual participant with the grand mean RDM minus that participant (Nili et al., 2014) (Figure 3A).

**Figure 3.**
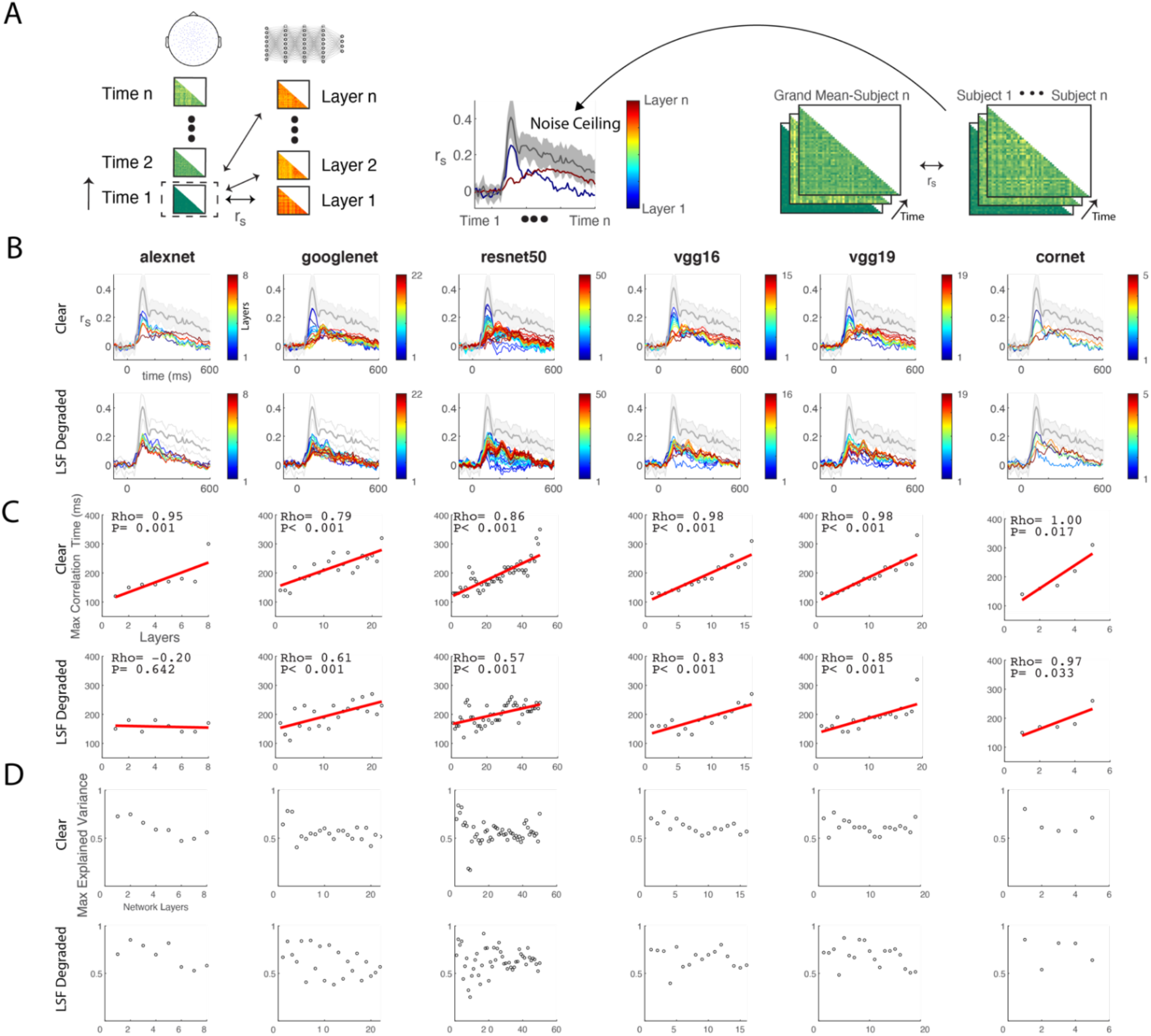
Temporal correspondence between MEG signals and Convolutional Neural Networks. (A) Schematic of the calculation to measure representational correspondence between MEG and CNNs. Spearman correlations were calculated iteratively in time between each participant’s MEG RDMs in 10ms increments from -100 to 600ms and across CNN RDMs derived from layer activations. Lower bound of the noise ceiling was calculated by iteratively correlating individual RDMs to the group mean RDM, excluding the individual RDM. Standard deviation is shown as shading around noise ceiling. (B) Time-resolved neural-CNN correspondence with x-axis as time in milliseconds and y-axis as Spearman rho. Color indicates CNN layer depth with blue representing shallow layers and red representing deep layers. (C) Top and bottom row show the time of maximum correspondence for each of the network layers with layers on x-axis and time in milliseconds on the y-axis. (D) Maximum explained variance calculated by neural-CNN correspondence divided by the lower bound of the noise ceiling for each CNN layer.

### Topographic Correspondence between Neural and Network RDMs

To assess how CNN-Brain correspondence changed as a function of sensor location, we constructed sensor by sensor RDMs using an electrode and its immediate surrounding neighbor sensors. As mentioned in the previous sections, we assessed CNN-Brain correspondence using Spearman correlations for each individual participant and then averaged the correlations across participants (Figure 4A). These results were tested for significance against zero correlation and corrected for multiple comparisons using FDR. To highlight the difference between clear and degraded images, we performed a pairwise test between conditions, correcting for multiple comparisons. For this analysis, we chose CORnet-S as it was found to be one of the most brain-like networks (Kubilius et al., 2018; Schrimpf, Kubilius, Hong, Majaj, Rajalingham, Issa, Kar, Bashivan, Prescott-Roy, Geiger, et al., 2018) consisting of only five layers (layers 1-5 are labeled V1, V2, V4, IT and Decoder) and ResNet-50 (included in the supplemental material) which was the largest net we tested.

**Figure 4.**
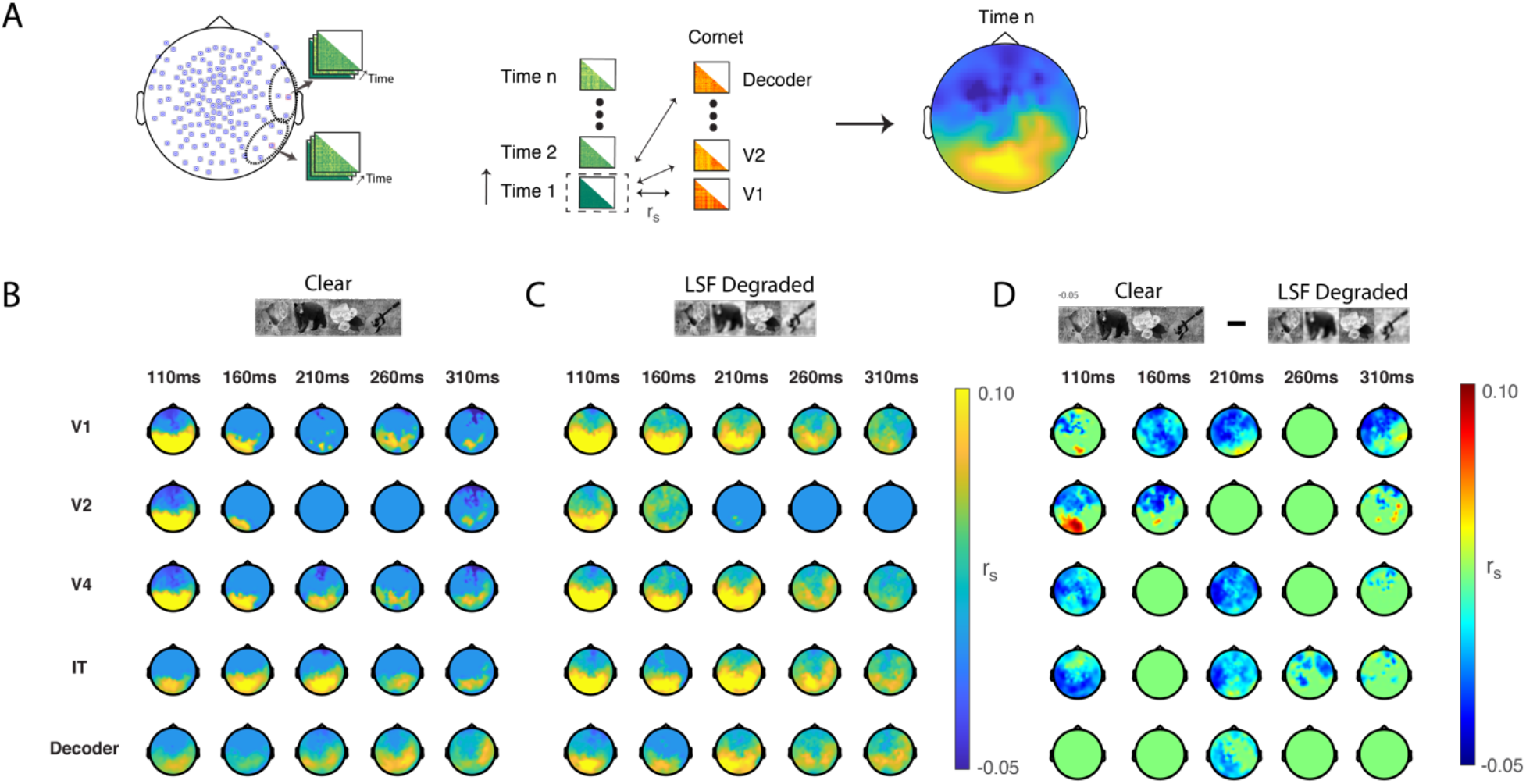
Topographic correspondence between MEG and CORnet-S. (A) Schematic of the searchlight procedure used to build electrode specific RDMs that can then be used to assess time resolved topographic correspondence between MEG and CORnet-S. (B-C) Topographic correspondence between all layers of CORnet-S at representative time periods to display how correspondence across MEG electrodes evolves in time. All correspondence is thresholded for significance using a Wilcoxon signed-rank test across participants against a null correlation, FDR corrected for multiple comparisons, q<0.05. (D) Differences in neural network correspondence between clear and degraded images at the same representative times, similarly thresholded for significance and corrected for multiple comparisons as in (B) and (C).

### Spectral Correspondence between Neural and Network RDMs

To capture spectral information, MEG signals were passed through a series of band-pass bidirectional Butterworth filters from 5 Hz to 45 Hz. We used a sliding window including the frequency of interest and 2 Hz above that frequency, such that 5 Hz represents 5-7 Hz, and 6 Hz represents 6-8 Hz, and so on and so forth. From the band-passed signals, we constructed frequency specific RDMs and then for each one of these frequencies measured the correspondence with CORnet-S RDMs and ResNet-50 RDMs (Figure 5A) for the same reasons described for the topographic correspondence. For ResNet-50, the RDMs were limited to one shallow, middle and deep layer; for CORnet-S, we included all the layers.

**Figure 5.**
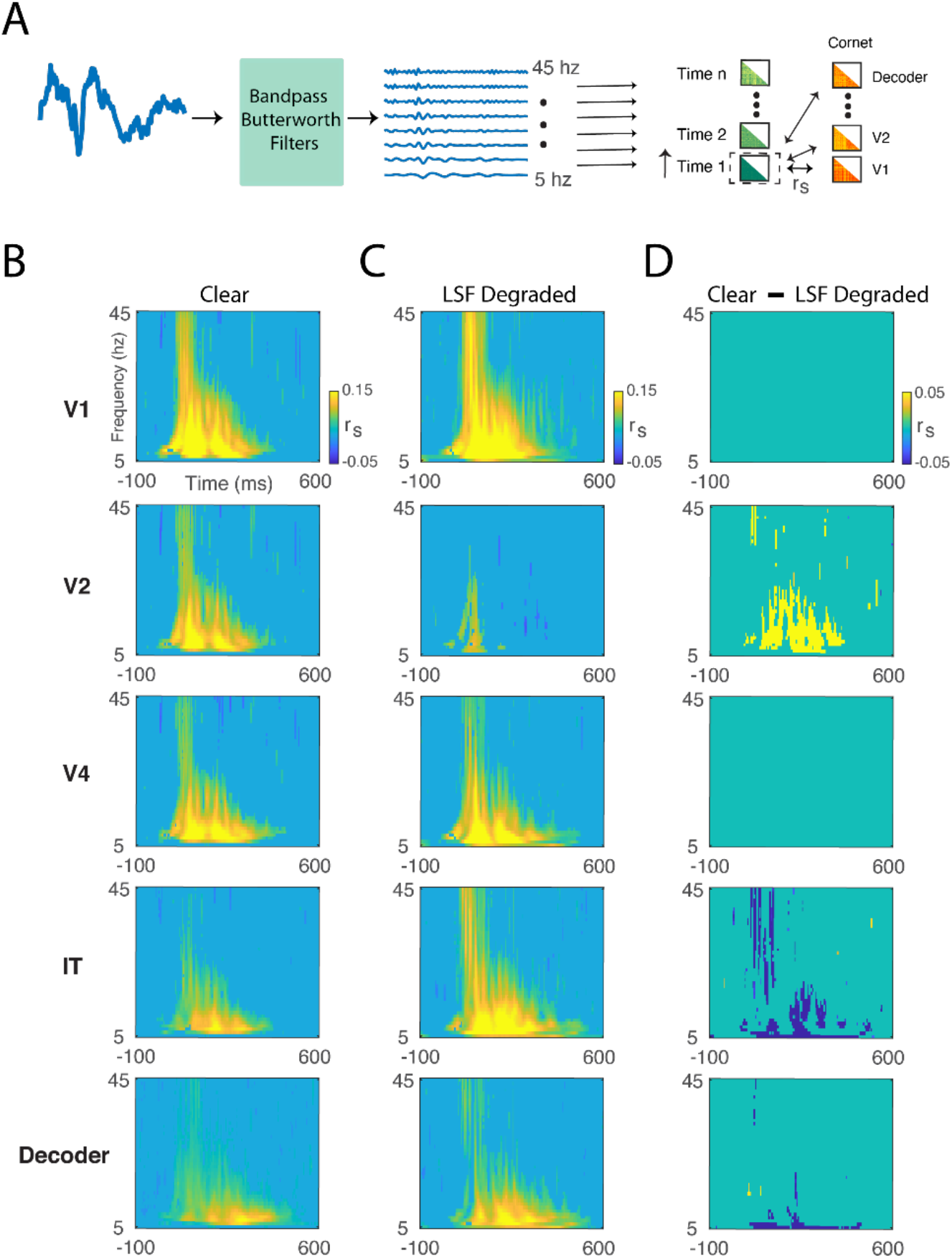
Time frequency correspondence between MEG and CORnet-S. (A) Schematic of time frequency analysis using 2nd order Butterworth filters to iteratively filter out frequency components. A 3 Hz sliding window, moving 1 Hz at a time until all frequency bands between 5-45 Hz were extracted. RDMs were then constructed at each frequency and correspondence between MEG frequency and CORnet-S was assessed (B-C) Time frequency correspondence between MEG signals and CORNet-S from shallow (top) to deep (bottom), thresholded for significance against 0 and corrected for multiple comparisons (q<0.05) for both clear (B) and degraded (C) images. (D) Significant difference between clear and degraded images for MEG-CORnet correspondence and corrected for multiple comparisons (q<0.05).

### Stylized image CNNs and CNN transfer learning

To test how CNN training, and specifically the features included within the images in the training set, affected CNN-Brain correspondence, we made use of a ResNet-50 architecture trained on a stylized ImageNet set (Geirhos et al., 2019), which we will refer to as “StyleNet”. The stylized images are the various images from ImageNet but with style transfer (Huang & Belongie, 2017) of textures from a diverse set of paintings (Figure 6A). For this network, the training parameters were as follows: 60 epochs with stochastic gradient decent, momentum term of 0.9, learning rate of 0.1 multiplied by 0.1 after 20 and 40 epochs, and a batch size of 256. In addition, we performed transfer learning on an AlexNet architecture, applying to ImageNet the low spatial frequency degradation that was used in the MEG experiment. Here, we used a degradation radius of 8 pixels on the cylinder in the sombrero convolution and applied this across all images. During transfer learning, we used a randomized subset of 250 of the 1000 image categories in ImageNet. The transfer learning parameters were as follows: 60 epochs with stochastic gradient decent, momentum term of 0.9, learning rate of 0.001, and batch size of 64. We then used these networks to measure the dynamics, CNN-Brain correspondence, and topographic CNN-Brain correspondence using the procedures described in the previous sections.

**Figure 6.**
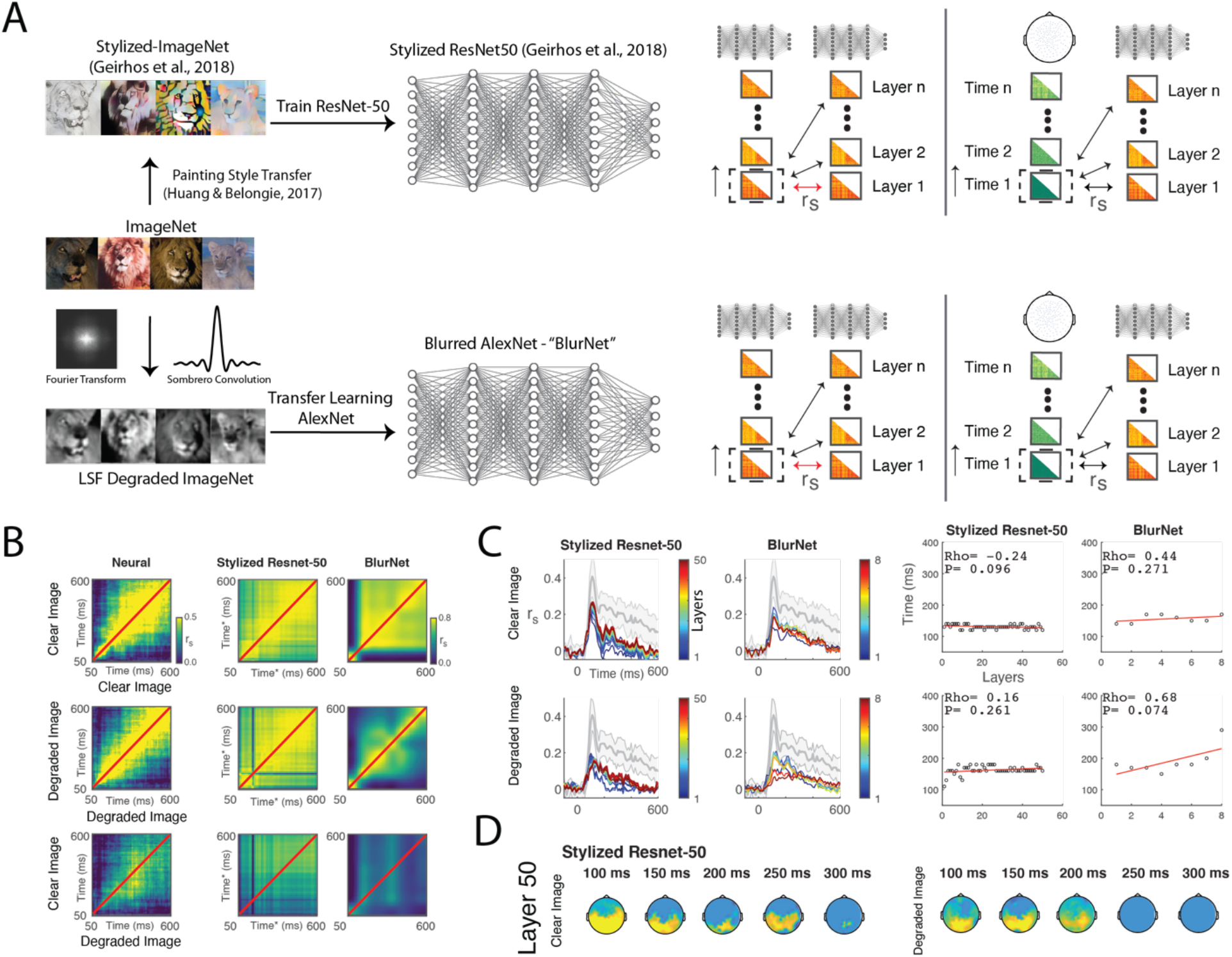
Assessing MEG-CNN correspondence with CNNs trained on stylized and low spatial frequency degraded images. (A) Schematic of procedures used to assess CNN trained on either a stylized version of ImageNet or on reduced ImageNet (250 categories) using LSF degraded image. Similar procedures as described in previous figures were then used to assess neural network correspondence. (B) Image processing dynamics as done in Figure 2 shown for StyleNet (Stylized ResNet-50) and BlurNet as well as previously shown neural dynamics for reference. (C) Temporal correspondence between modified CNNs and MEG (left panel) along with times of best correspondence for each layer. (D) Topographic correspondence for the last layers of StyleNet (Stylized-ResNet-50). See Supplemental Figure 4 for reference of ResNet-50 trained without image stylization.

## Results

### Difference between degraded low spatial frequency images and clear (i.e., unfiltered) images lateralizes to the right hemisphere

To determine when brain signals begin to diverge for low spatial frequency and clear, unfiltered images, we trained a classifier to distinguish between the two image types regardless of the specific exemplar. We found significant decoding onset at 50ms (Figure 1B), defined as at least two consecutive time points of significant decoding (Carlson et al., 2013). Decoding remained above chance throughout the stimulus period (500ms), peaking at 100ms post stimulus onset and extended to the end of the decoding window (100ms after stimulus offset). Using a searchlight analysis, we also measured topographic variation in the information regarding whether the image was clear or degraded. In the topographic maps, we found evidence of lateralization to the right hemisphere beginning at about 200ms and becoming more lateralized in time until 400ms. The lateralization of low spatial frequency information to the right hemisphere has been noted in previous studies (Flevaris & Robertson, 2016; Kauffmann, Ramanoël, & Peyrin, 2014; Schyns & Oliva, 1999). However, given that the difference between the low spatial frequency and the clear unfiltered image is the high frequency content, these results were somewhat surprising; high frequency information has been shown to lateralize to the left hemisphere (Flevaris & Robertson, 2016; Kauffmann et al., 2014; Schyns & Oliva, 1999). Thus, these results seem to suggest that the primary neural difference between the low spatial frequency and the unfiltered images are attributable to neural processing of low spatial frequencies.

### Relative responses to clear and low spatial frequency images are similar between neural networks and brains

We investigated the dynamics of how clear and degraded images were processed within brain signals and within CNNs by correlating RDMs for each respective model system (Figure 2A-B). In the correlation plots (Figure 2C-D), dark blue indicates low correlation between RDMs and bright yellow indicates higher correlations between RDMs. Qualitatively, we found similarities in the ways that both brain signals and CNNs process images (Figure 2C). While there were correlations in neighboring time points as well as between layers, there appeared to be a chain-like sequential processing of stimuli, such that the representational dynamics changed in time and across layers and no longer shared correlations to earlier times or layers. However, there were some notable differences from this general pattern. For example, CORnet-S had more shared similarity in shallow layers than deeper layers, a pattern that was in contrast to brain responses. For degraded images (Figure 2D), we found similar dynamics but found that there was relatively less correlation between neural signals and time as well as between CNN layers in architectures such as CORnet-S.

Of greatest relevance for our purposes, we computed the correlations between clear images and degraded images for both brain signals and CNN architectures (Figure 2E). Here, we found that degraded image information appears closely related to the information found in clear images at approximately 200ms. Moreover, the various CNN architectures embody this clear-degraded relationship to different degrees. To assess the similarity in the relative dynamics between CNN and brain signals for clear and degraded images, we calculated a similarity score by computing the squared Euclidean distance. We subtracted the total distance from one, such that higher scores indicate more similarity and lower scores indicate less similarity. The scores indicated that of all of the networks tested, VGG-19 had the most similar dynamic relationship between clear and degraded images to the brain. Overall, these results show that the dynamics and processing of clear and degraded images are similar between CNNs and brains.

### Degraded images lead to earlier CNN-Brain correspondence with deeper CNN layers

To directly assess correspondence between CNN activations and brain signals, we used a Spearman correlation to correlate the RDMs for each layer across CNNs in time. In general, we noted emergence of similar patterns across network architectures (Figure 3B and Supplemental Figure 3). For the clear images, there exist distinct peaks of correspondence for the shallow and deeper layers within the CNNs. In contrast, for the low spatial frequency degraded images, only a single peak was evident. We next quantified this observation by measuring the time at which the maximum correlation for each of the layers emerged (Figure 3C). First, we found that there was a positive correlation between layer depth and time across all architectures and across both types of image presentations with the exception of AlexNet with degraded images. Additionally, we found a steeper slope and higher degree of correlation for the clear images when compared with the degraded images across CNN architectures. We next measured the total amount of explained variance maximally achieved by each network architecture across each layer (Figure 3D). Using this approach, we found that the largest differences between clear and degraded images arose within the shallow layers.

In comparing across the various CNN architectures, we note some subtle differences between them. Deeper CNNs (i.e., those with more layers) as well as those that included recurrence showed sustained correlations later in the signal for deeper layers. For example, the last layer of CORnet-S had the highest explained variance (58.3%) at stimulus offset (500ms) for clear images. In comparison, the highest explained variance for ResNet-50 for clear images (51.0%) was seen 100 ms post stimulus offset (600ms). The highest explained variance regardless of layer depth or image presentation was found for ResNet-50 at 160ms in layer 17—a convolutional layer (res3c_branch2a). The high degree of explained variance (92%) was seen for degraded images. Collectively, the observed pattern of results imply that recurrence and deeper layers allow CNNs to be better models at higher stages of visual processing, agreeing with previous studies (Kietzmann, Spoerer, et al., 2019). Overall, we found that restricting an image to low spatial frequencies led to earlier CNN-Brain correspondence for the deeper layers when compared with clear images.

### Topographic CNN-Brain correspondence differs between clear and degraded images

Next, we quantified topographic correspondence between brain signals and CNNs by using a moving searchlight analysis to create electrode specific RDMs from the MEG signal. For this analysis, we limited the correlations to CORnet-S in our main analysis (Figure 4A) and ResNet-50 as a supplemental analysis. We chose CORnet-S due to the differences noted in the timings in the later layers in the previous section, its recurrence connections, and its relatively lower number of layers compared to other networks, allowing for easier visualization of the CNN-Brain correspondence. At 110ms, we find that CNN-Brain correspondence is primarily localized to the occipital MEG sensors across all CORnet-S layers. When we look at significant differences between the clear and degraded images (Figure 4D), we find that the correlation is significantly stronger for the clear images in layers V1 and V2 of CORnet-S. In comparison, the degraded images have stronger correspondence to frontal sensors, including sensors over orbitofrontal cortex. Over time, this pattern begins to change in such that CORnet-S layers V4 and IT show overall stronger correspondence with degraded images, including occipital sensors. Progressing forward in time, we find that the clear image correspondence stays fairly localized to visual cortex while the degraded image correspondence becomes more diffuse. This difference becomes most apparent at 210ms in nearly all layers except for layer V2. Progressing yet further in time, the CNN-Brain correspondence in later network layers is now lateralized to the right hemisphere and the differences between clear and degraded images become less apparent. However, topographic differences still exist in layer V1 with degraded images showing strong correspondence with frontal sensors and clear images showing some small localized increased correspondence in the right lateralized sensors. Together, these results show how the CNN-Brain correspondence in both time and across layers changes depending on whether participants and CNNs are processing clear images vs degraded images.

### Spectral CNN-Brain correspondence differs between clear and degraded images at different CORnet-S layers

Given the known existence of functional differences in information processing between frequency bands, such as gamma being more associated with feedforward processing and alpha and beta being more associated with feedback processing (Bastos et al., 2015; Belitski et al., 2008; Van Kerkoerle et al., 2014), we tested how correlations between brain signals and CNNs varied as a function of frequency. We again chose CORnet-S as the CNN for the reasons cited earlier. To extract frequency specific data, we used bidirectional Butterworth filters (Maier, Aura, & Leopold, 2011), capturing 3 Hz bands over the frequency range spanning 5 Hz to 45 Hz. After the results were tested for significance against zero correlation using a Wilcoxon signed rank test with FDR correction for multiple comparisons (Figure 5B and 5C), we found a general pattern emerge. In this pattern, early CORnet-S layers sharing broadband correspondence to brain signals, especially during the transient response for both clear and degraded images. Following this transient, frequency bands below 30 Hz captured the most correspondence between signals. Progressing into deeper CORnet-S layers, the correspondence was primarily localized to the lower frequency bands (<15hz).

In this frequency analysis, the difference between clear and degraded image correspondence showed a dissociation between network layers. The V2 layer in CORnet-S had higher correspondence in low frequency bands (< 30 Hz) for clear images than for degraded images. However, in deeper layers, specifically layer IT, degraded images had higher correspondence extending into the gamma range (30 - 45 Hz) for the transient peak. This advantage for degraded images was observed at the final decoder layers at low frequency bands (<30hz) during the sustained response, especially in the lowest frequency bands tested (5 Hz). In general, these findings support the notion that early CNN layers are more closely tuned to features that are present in brain signals of clear images but are missing from degraded images. In contrast, the degraded images, which still contain the conceptual aspects of the image, correspond more with the later layers of a neural network at low frequencies.

### Training CNNs with stylized images disrupts CNN-brain correspondence

Previous studies have shown that training CNNs with images of varying levels of abstraction shifts the focus of the CNN (such as StyleNet) more to shape rather than texture (Geirhos et al., 2019). Here, we tested the CNN-Brain correspondence for StyleNet and BlurNet. StyleNet is a ResNet-50 architecture trained on a stylized ImageNet image composed a wide variety of artistic styles. BlurNet is an AlexNet architecture that was trained using the same low spatial frequency manipulation used in the current study. As shown in figure 6B, we find that the dynamics in StyleNet are different than those seen in the brain, with each layer having shared representations with other layers. This pattern is seen for clear images as well as degraded images. Furthermore, when looking at the relationship between clear and degraded images, we find that clear-degraded generalization pattern in CNNs is different than the clear-degraded generalization pattern in the brain signals. Specifically, the RDMs for clear images in deeper layers correlate with degraded images across all layers for the CNNs but not for the brain. BlurNet showed similar dynamics for clear images as StyleNet with widely shared representations following the initial layers. Interestingly with the degraded images, there was a more chain-like dynamic as observed in the CNNs in Figure 2. However, the clear-degraded generalization pattern was again different from what was observed in the brain signals.

When looking at the direct CNN-Brain correspondence, we see that all of the layers correspond to early times within the MEG signal (Figure 6C). This result most likely reflects the shared correlation between the layers shown in Figure 6B. For clear images, the explained variance at later times dropped from what was found in the ResNet-50 architecture with StyleNet explained variances at layer 50 of 0.5% and -8.0% at 500ms and 600ms. In comparison, the last layer in ImageNet-trained ResNet-50 yielded explained variance of 49.9% and 51.0%. For BlurNet, explained variance to clear images was 10.6% and 11.9% at 500ms and 600ms while AlexNet had explained variances of 36.3% and 21.9%.

For degraded images, the CNN-Brain correspondence decreased for StyleNet but improved for BlurNet. StyleNet had explained variance of -15.5% and 12.5% at 500ms and 600ms while the comparable values for ResNet-50 at these times was 23.3% and 23.0%. The last layer of BlurNet had explained variance of 31.2% and 25.1% at 500ms and 600ms while the last layer of AlexNet had explained variance of 14.8% and -1.7% at those times. However, there was no longer a direct linear relationship in time and within layers for either StyleNet or BlurNet for degraded images. Lastly, the late layers for both clear and degraded images in StyleNet localized to occipital sensors (Figure 6D) across time points. Overall, these results show that training a neural network with stylized images leads to poor correspondence with brain responses, especially for signals in the later portions of the evoked MEG response to stimuli; the notable exception to this generalization are results from BlurNet on degraded images.

## Discussion

In this study, we investigated the effects of image perturbations, notably LSF blurring, on how well CNNs modeled dynamic brain signals by measuring layer-by-layer correspondence between CNNs and the time resolved MEG signal. The major finding of the study is that CNN-Brain correspondence emerged earlier in time when images were degraded than when they were clear. When comparing brain activity associated with viewing clear vs. degraded images, we found that decoding was lateralized to MEG sensors in the right hemisphere, the brain hemisphere that preferentially processes low spatial frequency visual information (Flevaris & Robertson, 2016; Kauffmann et al., 2014; Schyns & Oliva, 1999). These results suggest that the earlier CNN-Brain correspondence to degraded images is primarily driven by differences in how the brain processes low spatial frequencies. The absence of high spatial frequency content in the blurred images effectively boosted the impact of the low spatial frequency information on brain activity, perhaps through what Bar (2021) refers to as “initial guesses” about what one is viewing that is signaled via feedback from higher brain areas. The CNN-Brain topographic results further fit within a broader coarse-to-fine theoretical framework (Bar, 2003b, 2021; Goddard, Carlson, Dermody, & Woolgar, 2016; Kauffmann et al., 2015; Lu et al., 2018) in which we find correspondence between early visual sensory areas and shallow CNN layers early in time for clear unfiltered images while degraded low spatial frequency images have stronger correspondence to deeper CNN layers and MEG sensors in frontal areas soon after stimulus presentation.

Our findings are at odds with recent fMRI results pointing to shared Brain-CNN correspondence within low level visual areas but not high level visual areas, and particularly decreased correspondence with degraded images (Xu & Vaziri-Pashkam, 2021). We believe these apparent contradictions are attributable, at least in part, to the temporal fine structure that can be resolved in MEG signals but not in fMRI BOLD signal. For example, Xu and Vaziri-Pashkam (2021) found that ResNet-50 was one of the only CNNs that had shared correspondence with higher level visual areas. Similarly, we found that ResNet-50 accounted for the greatest variance in later times of the MEG evoked response for clear images (i.e.,100ms following stimulus offset). However, by using the time-resolved MEG signal, we also found that earlier correspondence for degraded images was localized in MEG parietal and frontal sensors. Thus, we conjecture that fMRI studies are unable to resolve this aspect of CNN-Brain correspondence owing to the sluggishness of the BOLD response. In turn, this suggests that the fMRI signal is likely to be unable to register signals associated with recurrent dynamics, signals that are best captured with recurrent CNNs. Indeed, our study showed that CORnet-S improved late brain-fMRI correspondence compared with other CNNs that did not have recurrent connections, in agreement with previous work (Kietzmann, Spoerer, et al., 2019).

Beyond demonstrating that aspects of CNN-Brain correspondence may be obscured within the sluggish BOLD signals measured using fMRI, MEG studies reveal a key temporal correspondence between brain signals and CNN layers: early brain signals correspond to shallow CNN layers and late brain signals correspond to deep CNN layers (Cichy et al., 2016; Greene & Hansen, 2018; Kietzmann, Spoerer, et al., 2019; Kong et al., 2020; Seeliger et al., 2018). Thus, information is lost if we do not account for the time varying signals that the brain uses (Carlson et al., 2013; Cichy et al., 2014) when measuring correspondence to object processing in the layers of a CNN. In our study, we found such temporal correspondence but further leveraged this relationship and specifically probed the dynamics in time and between CNN layers by generalizing RDMs in time as well as across layers. We found that not only were there shared dynamics in processing clear and degraded images, but also similarities in the way that CNNs and brains respond to image perturbations. By using dynamics to gauge for similarity in processing dynamics, we were able to learn another important lesson: when CNNs are trained using stylized image sets (Geirhos et al., 2019) or degraded image sets, they no longer share similar processing dynamics as the brain, despite explaining comparable variance during the peak of the MEG signal. From this, we put forth that when modifying training sets to build CNNs that can serve as better models of the brain (Mehrer et al., 2021), measuring dynamics to image perturbations may serve as an effective metric to index CNN-brain correspondence.

The dynamic MEG signal also allows one to probe how correspondence between brains and CNNs may change as a function of brain oscillations. We found correspondence was strongest between the V2 layers of CORnet-S and MEG signals for clear images during the sustained response and predominated in the alpha/beta range to lower theta frequency bands. However, this pattern reversed with higher correspondence in deeper CORnet-S layers for degraded images in the gamma band during the transient and theta band during the sustained response. These findings are consistent with earlier work showing that low spatial frequency image information is preferentially carried in gamma bands while higher frequency image information is preferentially carried in alpha bands (Bar, Kassam, Ghuman, Boyshan, et al., 2006; Flevaris & Robertson, 2016; Fründ, Busch, Körner, Schadow, & Herrmann, 2007). Additionally, gamma band oscillations have also been linked with magnocellular and dorsal stream activity (Merigan & Maunsell, 1993; Tootell et al., 1988), which ostensibly carry the coarse information in the coarse-to-fine processing framework (Bar, Kassam, Ghuman, Boshuan, et al., 2006). The differences found between frequency bands in MEG signals provides motivation to further investigate the correspondence in laminar and direct local field potential (LFP) recordings, which have shown rich frequency specific LFP differences in feedforward and feedback processes within localized circuits (Bastos et al., 2012; Bastos et al., 2015; Maier et al., 2011; Mineault, Zanos, & Pack, 2013; Van Kerkoerle et al., 2014). For such studies, there are a number of potential targets including the distinct magno- and parvocellular layers in LGN (Poltoratski, Ling, McCormack, & Tong, 2017; Tootell et al., 1988), V1 layers where spatial frequency continues to be dissociated between layers 4Cb and 4Ca respectively (Tootell et al., 1988), as well as area V4 which contains separate low spatial frequency and high spatial frequency domains (Lu et al., 2018).

The general tendency we observed for deep CNN layers to show higher correspondence with degraded images earlier in time may point to categorical commonalities between CNNs and brains that are largely missing at the exemplar level (Rajalingham et al., 2018). Since low spatial frequency images prompt the processing of visual images at the superordinate level, which defines category wide attributes (Ashtiani, Kheradpisheh, Masquelier, & Ganjtabesh, 2017), individual CNN-Brain correspondence may become higher as the exemplar become less distinguishable and the images are reduced to possible membership in broad categories. In addition, correspondence could be improved through modifications to CNNs that create more stable exemplar representations. For example, a recent study found that exemplar representations vary between network initializations (Mehrer, Spoerer, Kriegeskorte, & Kietzmann, 2020), and that averaging across several different initializations can improve CNN representations. Alternatively, CNNs trained on datasets that include object categories that are more relevant to humans rather than those comprising ImageNet, which includes an overemphasis on categories such as dog breeds, could also provide more brain-like exemplar representations (Mehrer et al., 2021). Finally, another potential avenue to explore are CNNs that have been trained on sets of low spatial frequency images with decreasing degrees of blur, thus simulating visual development in infants. CNNs trained in that way have shown better performance than CNNs trained on unblurred images from the outset, leading to the speculation that graded training makes the CNN more brain-like (Avbersek, Zeman, & Op de Beeck, 2021). Probing different modifications to CNN training paradigms will be essential in testing how the image statistics in trainings affect the Brain-CNN correspondence across a number of different image perturbations and differing levels of occlusion (Rajaei, Mohsenzadeh, Ebrahimpour, & Khaligh-Razavi, 2019; Schrimpf, Kubilius, Hong, Majaj, Rajalingham, Issa, Kar, Bashivan, Prescott-Roy, Geiger, et al., 2018).

## Conclusion

In conclusion, we have provided evidence of earlier correspondence between brains and deep CNN layers in degraded images that support the coarse-to-fine conceptual framework of visual image processing. In addition, we have provided a rich methodological framework by introducing a number of analyses that can be used to assess the dynamics of CNNs and compare these with brain activity across the dimensions of space, time, and frequency. This framework can be extended to include a number of image perturbations as we test the limits of CNN-brain correspondence with CNNs that are purposefully created to be more brain-like (Kubilius et al., 2018) or those that inadvertently become so (Schrimpf, Kubilius, Hong, Majaj, Rajalingham, Issa, Kar, Bashivan, Prescott-Roy, Geiger, et al., 2018). Finally, there are a number of potentially revealing experimental manipulations that could enhance efforts to examine possible CNN-brain correspondence. Those include manipulations of stimulus duration (Grootswagers, Robinson, & Carlson, 2019), creation of visual stimuli comprising object textures devoid of explicit shapes (Grootswagers, Robinson, Shatek, & Carlson, 2019; Long, Yu, & Konkle, 2018), visual images that are accompanied by congruent or incongruent sounds (Tovar, Murray, & Wallace, 2020), and creation of hybrid stimuli consisting of conflicting low spatial frequency and high spatial frequency information (Schyns & Oliva, 1999). These kinds of manipulations, together with expanded CNN architectures and training sets, will push the boundaries of understanding of the potential correspondence between brains and CNNs.

## Acknowledgements

This work was supported by the National Institute of General Medical Sciences of the National Institutes of Health (Grant T32-GM-007347). RB and OC were supported by research funds associated with RB’s Vanderbilt University Centennial Professorship.

## Supplemental Figures

**Supplemental for Figure 1.**
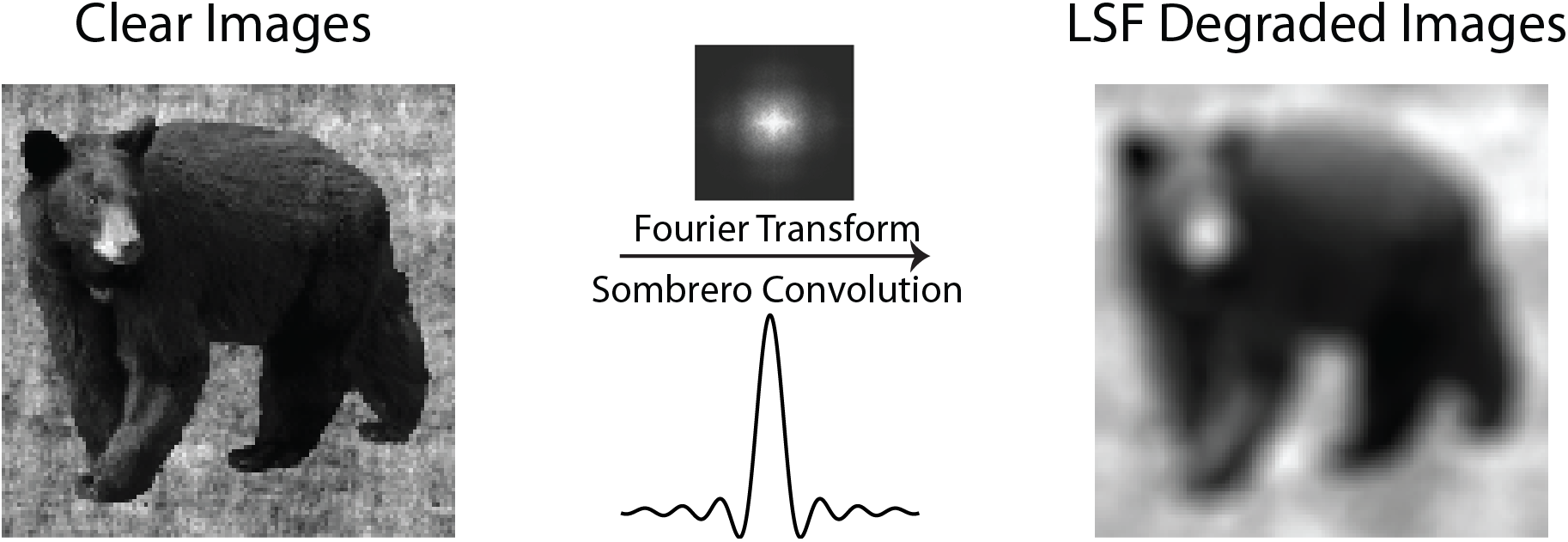
Example LSF degradation of one of the exemplars

**Supplemental for Figure 3.**
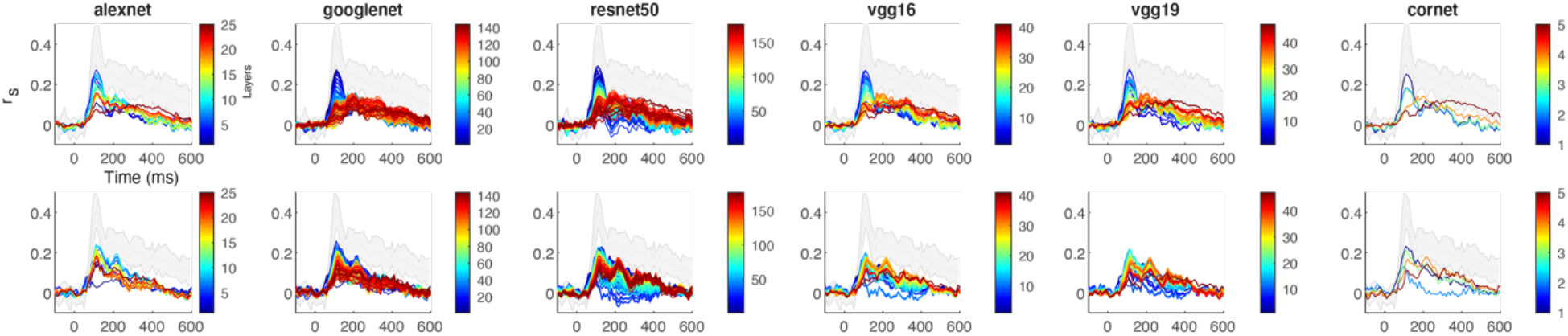
Supplemental correspondence between MEG and CNNs using all operations including pooling, ReLU, and normalization

**Supplemental for Figure 4.**
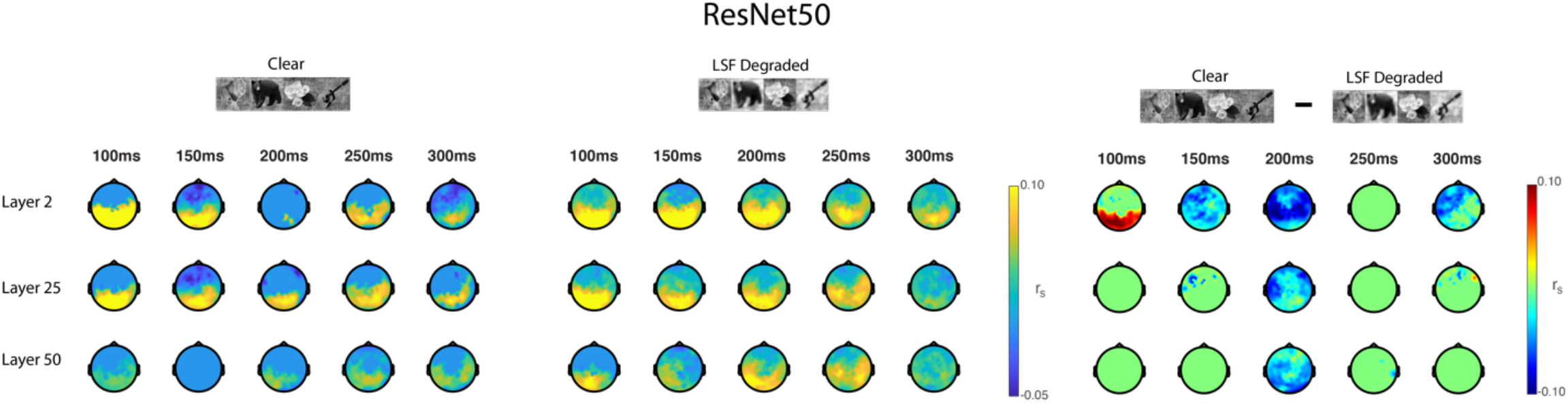
Supplemental topographic correspondence between MEG and ResNet50 that largely complement the findings found using CORnet-S

**Supplemental 1 for Figure 5.**
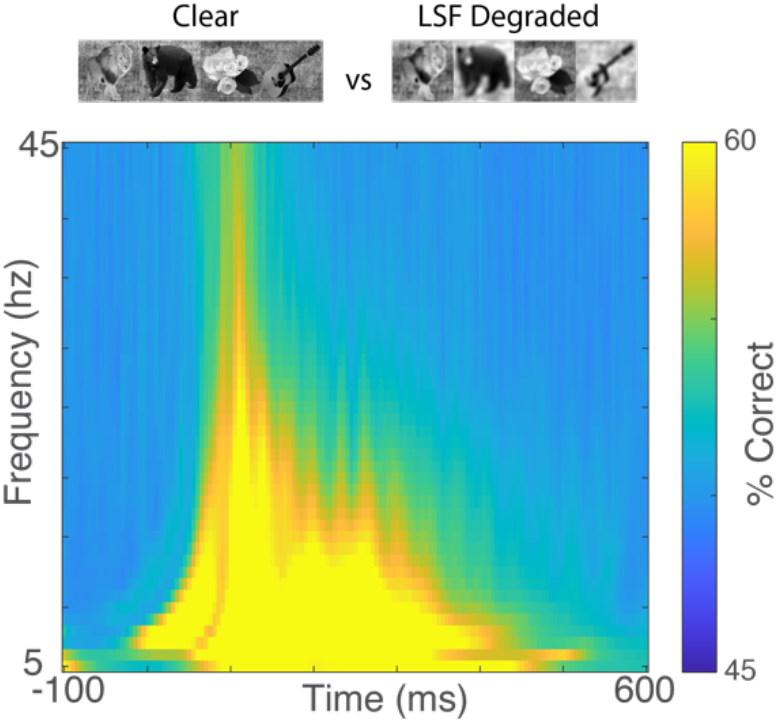
Time frequency decoding between clear and degraded images, showing the spectrotemporal profile differentiating between the images.

**Supplemental 2 for Figure 5.**
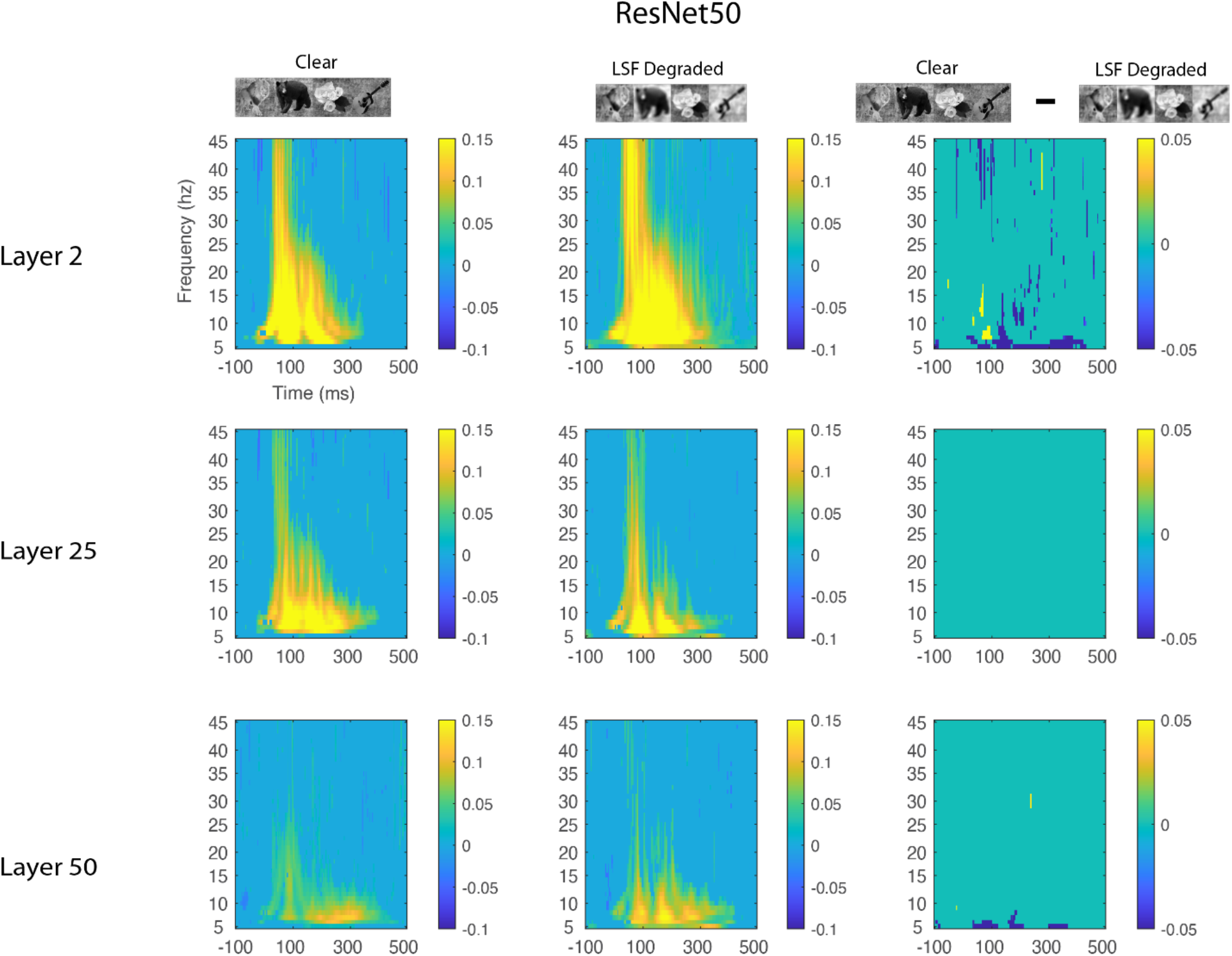
Supplemental spectrotemporal correspondence between MEG and ResNet50 that shows some differences between CORnet-S and ResNet50. Namely, that there is higher correspondence between degraded images and ResNet50. However, note that these are not all of the layers in ResNet50, just representative of shallow, middle, and deep layers.

